# Validation and characterisation of a wheat GENIE3 network using an independent RNA-Seq dataset

**DOI:** 10.1101/684183

**Authors:** Sophie A. Harrington, Anna E. Backhaus, Ajit Singh, Keywan Hassani-Pak, Cristobal Uauy

## Abstract

Gene regulatory networks are powerful tools which facilitate hypothesis generation and candidate gene discovery. However, the extent to which the network predictions are biologically relevant is often unclear. Recently, as part of an analysis of the RefSeqv1.0 wheat transcriptome, a GENIE3 network which predicted targets of wheat transcription factors was produced. Here we have used an independent and publicly-available RNA-Seq dataset to validate the predictions of the wheat GENIE3 network for the senescence-regulating transcription factor *NAM-A1* (TraesCS6A02G108300). We re-analysed the RNA-Seq data against the RefSeqv1.0 genome and identified a *de novo* set of differentially expressed genes (DEGs) between the wild-type and *nam-a1* mutant which recapitulated the known role of *NAM-A1* in senescence and nutrient remobilisation. We found that the GENIE3-predicted target genes of *NAM-A1* overlap significantly with the *de novo* DEGs, more than would be expected for a random transcription factor. Based on high levels of overlap between GENIE3-predicted target genes and the *de novo* DEGs, we also identified a set of candidate senescence regulators. We then explored genome-wide trends in the network related to polyploidy and homoeolog expression levels and found that only homoeologous transcription factors are likely to share predicted targets in common. However, homoeologs in dynamic triads, i.e. with higher variation in homoeolog expression levels across tissues, are less likely to share predicted targets than stable triads. This suggests that homoeologs in dynamic triads are more likely to act on distinct pathways. This work demonstrates that the wheat GENIE3 network can provide biologically-relevant predictions of transcription factor targets, which can be used for candidate gene prediction and for global analyses of transcription factor function. The GENIE3 network has now been integrated into the KnetMiner web application, facilitating its use in future studies.

## Introduction

Transcriptional regulation of gene expression is fundamental to all biological processes. Increasingly, this is studied using large-scale datasets obtained from RNA-Sequencing (RNA-Seq) experiments across multiple tissues, genotypes, treatments, and timepoints. As library preparation and sequencing costs fall, more and more RNA-Seq datasets are being published, providing a wealth of transcriptional information. These datasets are then available for integration into large-scale gene regulatory networks covering various biological conditions. In wheat (*Triticum aestivum*), genomic and transcriptomic analysis has historically been hampered by its large, repetitive polyploid genome [1]. More recently, a high-quality genome and gene annotation has facilitated transcriptomics work in wheat [1, 2]. This has allowed the use of substantial RNA-Seq datasets to build gene regulatory networks and predict transcription factors involved in complex processes such as senescence [3] and grain development [4, 5]. However, these studies typically use bespoke RNA-Seq datasets to generate the regulatory networks, rather than exploiting publicly-available data.

Combining the data across multiple RNA-Seq studies allows new hypothesis generation beyond those possible in the initial individual publications. A key difficulty, however, in using publicly available datasets is the lack of consistency in metadata (e.g. relating to the experimental conditions, tissue sampling, developmental stage). Without clear labelling of sampling conditions, it is difficult, if not impossible, to compare RNA-Seq datasets that originated from different studies. While international efforts to standardise experimental data exist [6, 7], often these are not followed by researchers when publishing. To address this, a large effort was recently taken in polyploid wheat to integrate publicly-available RNA-Seq datasets into a common database (expVIP) [8]. This platform uses common metadata terms, manually annotated by the curators, to allow comparison between RNA-Seq datasets from different studies. This database has been updated with new wheat gene annotations, most recently following the release of the wheat reference genome (RefSeqv1.0) [2].

These large curated sets of transcriptome data can now be mined to build new gene regulatory networks covering many biological processes in wheat. Using 850 RNA-Seq datasets combined from multiple independent studies, gene co-expression networks for root, leaf, spike, and grain tissues, as well as abiotic and biotic stresses, were developed [2]. This same study also generated a network of predicting transcription factor - target relationships using the 850 independent RNA-Seq samples from a wide range of developmental, tissue, genotypes and stress conditions [2]. This network was created with the GENIE3 algorithm, which uses a Random Forests approach to predict the strength of putative regulatory links between a target gene and the expression pattern of input genes (i.e. transcription factors) [9]. The program produces a ranked list output of each pairwise comparison ranked from the most confident to the least confident regulatory connection. GENIE3 was able to recapitulate known genetic regulatory networks in *Escherichia coli* when first tested. Since its introduction, the GENIE3 algorithm has been used to identify tissue-specific gene regulatory networks in maize [10] and key regulatory genes in glaucoma [11], as well as to study the drought response in sunflower [12]. Previous studies have integrated the GENIE3 network predictions with ChIP-Seq and other proteomic and transcriptomic data and found that the GENIE3 predictions do correspond with independent biological datasets [10, 13].

Here we have conducted a series of analyses to investigate whether the GENIE3 network provides biologically-relevant information in polyploid wheat. As a first case study, we re-analysed the RNA-Seq datasets from Pearce *et al.* [14] which examined gene expression of the NAC transcription factor *NAM-A1*. This transcription factor is known to affect monocarpic senescence and nutrient remobilisation in polyploid wheat, affecting gene expression even before visual signs of senescence can be observed (e.g. 12 days after anthesis in flag leaf) [14, 15]. We compared the differentially expressed genes between wild-type and *nam-a1* mutant lines with the GENIE3 predicted targets of the *NAM-A1* transcription factor [14]. This publicly-available RNA-Seq data was not used in the generation of the GENIE3 network, thus serving as an independent dataset for validation purposes. We then explored the GENIE3 network for genome-wide trends relating to polyploidy and investigated the putative functions of targets for wheat transcription factors. Finally, we integrated the GENIE3 network into the KnetMiner web application [16] to facilitate exploration of the data within a wider context.

## Methods

### GENIE3

The GENIE3 network was previously published in [2] and made available at https://doi.org/10.5447/ipk/2018/7. In brief, it utilised a set of 850 publicly-available RNA-Seq samples in a Random Forests approach to predict targets of wheat 3,384 transcription factors [9]. The top one million connections in the network were used for all analyses in the paper, consistent with previous studies [2, 10]. Target genes of particular transcription factors were extracted from the network as in “Genie3_Statistics_SharedRatios_RNASeqDEGs_Figure1.Rmd” at https://github.com/Uauy-Lab/GENIE3_scripts/.

### RNA-Seq analysis

#### Mapping

Publicly-available reads from [14] were downloaded from https://www.ncbi.nlm.nih.gov/geo/query/acc.cgi?acc=GSE60635. Reads from the wild-type and *nam-a1* (*gpc-a1*) mutant lines from 12 and 22 days after anthesis (DAA) were pseudoaligned with kallisto (v 0.43.1) [17] against the v1.1 annotation from the RefSeq genome v1.0 [1] using standard settings for single reads (--single -b 30 -l 200 -s 20) (Supplementary Table 1). Only the A and B genomes of RefSeqv1.0 were used for the pseudoalignment, as the raw reads were derived from tetraploid *cv*. Kronos plants.

#### Differential Expression Analysis

Gene expression levels (transcript per million, TPM) were determined using the R package Sleuth [18] using the default settings for the Wald test (sleuth_wt; v 0.30.0). We compared the expression of genes between the wild-type and *nam-a1* mutant samples at 12 and 22 DAA. We used the cut-off of *q* < 0.05 to identify differentially expressed genes, where *q* is the *p*-value adjusted for false discovery rate using the Benjamini-Hochberg correction. The list of differentially expressed genes for each timepoint is reported in Supplementary Table 2.

#### Methods for ID conversion and comparison

The genes and contigs identified as differentially expressed in the original [14] study were converted to RefSeqv1.1 gene models where possible using BLASTn [19]. Briefly, the differentially expressed sequences were extracted from the IWGSC CSS genome [20] and compared with BLASTn (v 2.2.3; - num_alignments 1 -outfmt 6) against the RefSeq v1.1 transcriptome (including both high and low confidence gene models). The BLAST hit with the greatest percent identity to the original CSS sequence was assigned as the equivalent RefSeqv1.1 gene model. The scripts used for this analysis are found at https://github.com/Uauy-Lab/GENIE3_scripts, ExtractCDS_fromPearceDEGs.py and BLAST_Pearce_cds_against_HC_and_LC.sh.

### Comparison of the Differentially Expressed Genes with GENIE3

#### Calculation of Shared Ratios

We calculated the level of overlap or shared genes between different transcription factors or datasets as follows:

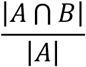

Where A and B are sets of genes, and |A| < |B|.

For example, given two sets of genes A and B, where A contains 5 genes and B contains 10 genes, if they share two genes in common the shared ratio is 2/5, or 0.4.

This calculation was carried out as implemented in R scripts available at https://github.com/Uauy-Lab/GENIE3_scripts. It was used to create the distribution of shared targets between transcription factors and the differentially expressed genes, as well as between the predicted targets of any two transcription factors.

#### Distributions of Shared Ratios

Initially, we analysed the shared ratios between transcription factors in the GENIE3 network and the set of differentially expressed genes obtained from the re-analysed *NAM-A1* RNA-Seq data. The target genes of 1000 randomly selected transcription factors were compared against the differentially expressed genes at 12 and 22 DAA to obtain the distribution of shared ratios (Figure 1B). This calculation was also carried out individually for the targets of *NAM-A1* against both timepoints, as implemented in https://github.com/Uauy-Lab/GENIE3_scripts/Genie3_Statistics_SharedRatios_RNASeqDEGs_Figure1.Rmd.

**Figure 1:**
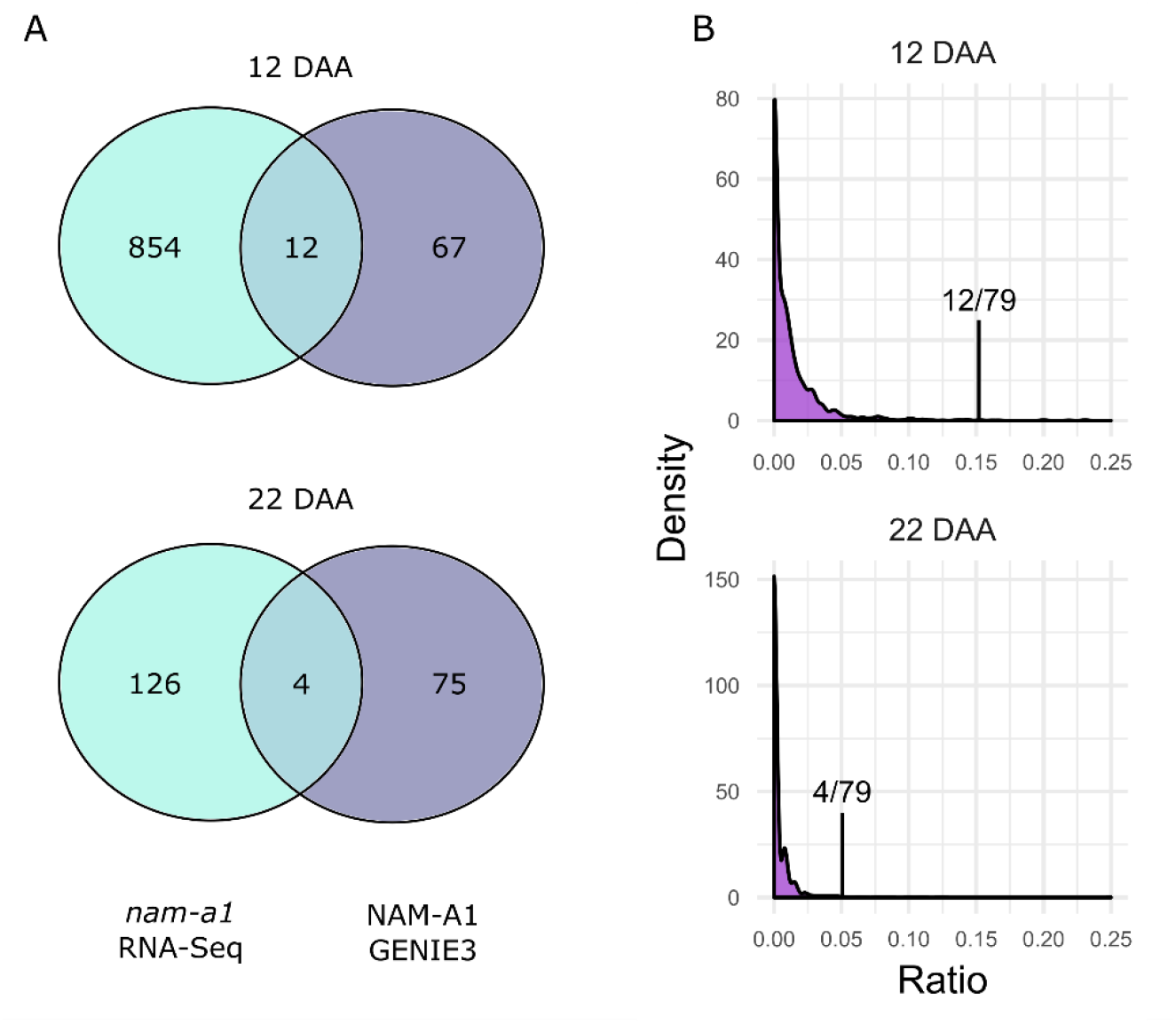
GENIE3 predicts targets of NAM-A1 that overlap with genes differentially expressed in *nam-a1* mutantsn. (**A**) More overlapping genes are identified at 12 DAA (15.2% of the GENIE3 targets) than at 22 DAA (5.1%). (**B**) Most transcription factors share very few predicted targets with the *nam-a1* differentially expressed genes, with a distribution skewed strongly towards 0. At both time points, the predicted targets of NAM-A1 overlap significantly more with the *nam-a1* DEG than would be expected by chance (Sign Test, p < 0.001). Note that the x-axis is capped at 0.25, to emphasize the skew of the distributions towards zero. “Ratio” refers to the shared ratio of targets between the DEGs and the GENIE3 transcription factors.

Following this, the shared ratio was calculated for 1000 randomly selected pairs of transcription factors from the GENIE3 network (implemented in https://github.com/Uauy-Lab/GENIE3_scripts/Genie3_Statistics_SharedRatios_AllTFs_AllTFFamilies_Fig3_SuppFig2.Rmd) (see Figure 3A). The same analysis was carried out for individual transcription factor super-families, based on the family assignments from https://opendata.earlham.ac.uk/wheat/under_license/toronto/Ramirez-Gonzalez_etal_2018-06025-Transcriptome-Landscape/data/data_tables/transcription_factors_to_use_high_confidence.csv [2]. All pairs selected were unique, and where a transcription factor family was not large enough to contain 1000 unique pairs, the maximum number of unique pairs was sampled (e.g. in the family CCAAT_HAP3, N=3 and thus the number of unique pairs sampled was 6). This calculation was also carried out for all homoeolog pairs, where triads were classified as in https://opendata.earlham.ac.uk/wheat/under_license/toronto/Ramirez-Gonzalez_etal_2018-06025-Transcriptome-Landscape/data/TablesForExploration/Triads.rds [2]. The subset used for this analysis only included syntenic 1:1:1 triads (see [2] for definition), resulting in a total of 708 triads and 2,124 individual genes. This was implemented in https://github.com/Uauy-Lab/GENIE3_scripts/Genie3_Statistics_SharedRatios_Homoeologs_MovementCategories_Figure4_SuppFig3.Rmd.

#### Movement Ratios

The shared ratios of homoeologous pairs were also distinguished by movement classifications, as defined previously [2]. In brief, triads were classified into three categories (“Dynamic”, “Middle 80”, and “Stable”) based on variation in their homoeolog expression bias across tissues. Dynamic triads show more variation in relative homoeolog expression levels across tissues than stable triads. The assignment of triads to each category is found here: https://opendata.earlham.ac.uk/wheat/under_license/toronto/Ramirez-Gonzalez_etal_2018-06025-Transcriptome-Landscape/data/Triad_Subsets_Movement/. Triads in the “HC_CS_no_stress_movement_top_10pc.txt” file were defined as Dynamic, the “HC_CS_no_stress_movement_middle_80pc.txt” as Mid 80, and the “HC_CS_no_stress_movement_low_10pc.txt” as Stable. This analysis was implemented in “Genie3_Statistics_SharedRatios_Homoeologs_MovementCategories_Figure4_SuppFig3.Rmd” at https://github.com/Uauy-Lab/GENIE3_scripts.

#### Developmental Expression Datasets

Public datasets were used for the expression analysis in Figure 2. The developmental time course was first published in [2], from the spring wheat *cv.* Azhurnaya. This dataset was included in the generation of the GENIE3 network. The senescence-specific time course was first published in [3], from the spring wheat cultivar Bobwhite, and was not included in the GENIE3 network.

**Figure 2:**
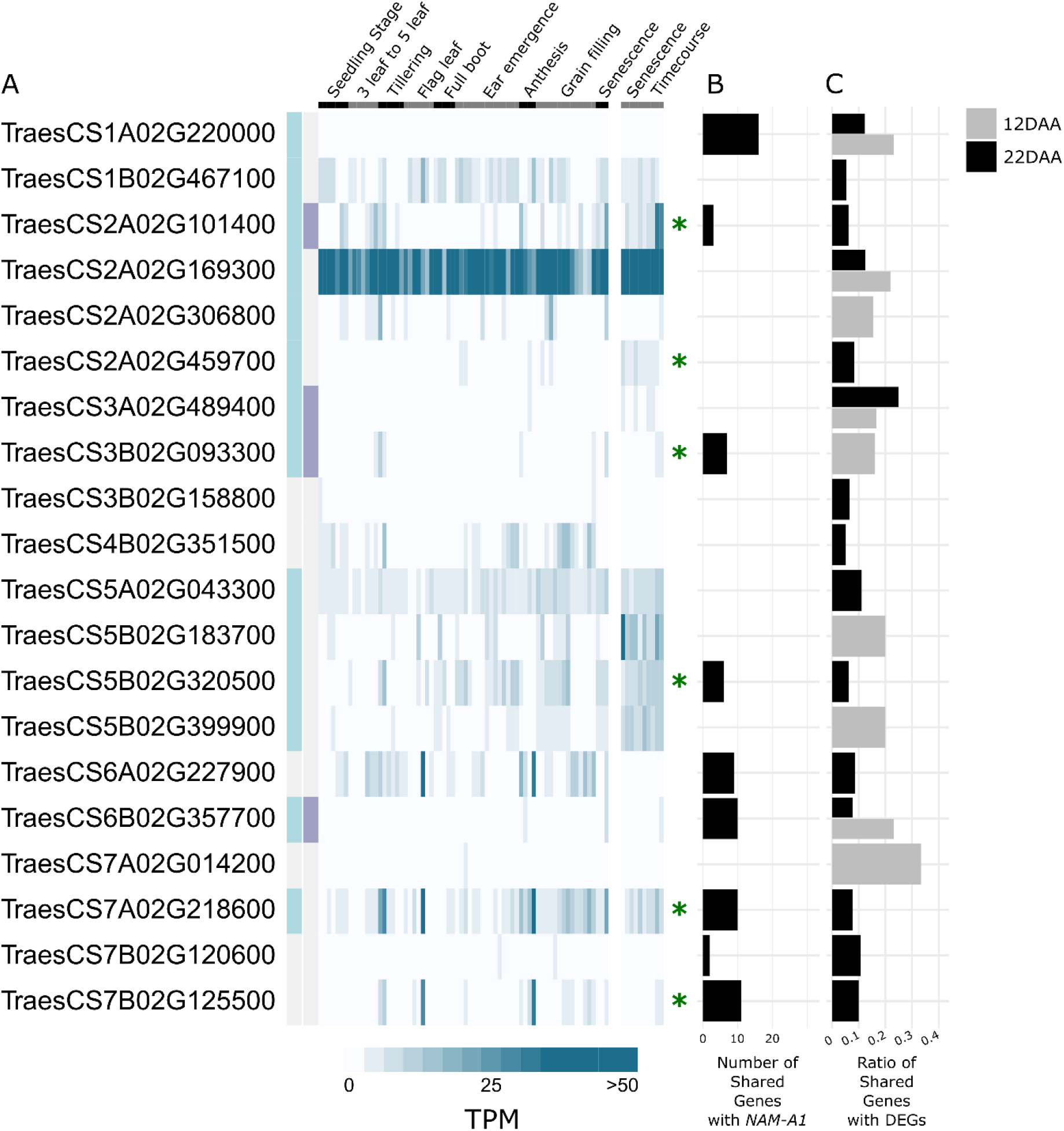
Candidate senescence regulators. Twenty transcription factors were identified which had a higher shared ratio between the GENIE3-predicted targets and the RNA-Seq DEGs than *NAM-A1* itself. (**A**) Their expression pattern is shown across a developmental time course [2] and a senescence-specific time course [3]. The TPM reported is averaged across three samples of the same tissue and timepoint from each dataset. Genes upregulated in senescence are highlighted with a light blue box, and those with a greater than 2-fold increase are highlighted with a purple box (left side). Genes present in the list of 341 candidate transcription factors based on the senescence time course in Borrill *et al.* 2019 are indicated with a green asterisk [3]. (**B**) The number of targets shared between the transcription factors and *NAM-A1*. (**C**) The shared ratio for each gene against the 12 DAA (grey) and 22 DAA (black) DEGs. Note that genes which had a higher shared ratio at both 12 and 22 DAA are shown with split bars.

### Gene Ontology (GO) term analysis

GO-term enrichment analysis was carried out as previously described in [3], using the GOSeq (v 1.34.1) package in R [21].

### Data visualisation, manipulation, and statistical analyses

Graphs were made in R, principally using the ggplot2 (v 3.1.1)[22] and ggpubr (v 0.2)[23] packages as well as the “aheatmap” function of the NMF package (v 0.21.0)[24]. Networks in Figures 3B and C were visualised using Cytoscape (v 3.7.1) [25]. Data manipulation was also carried out in R, using the packages dplyr (v 0.8.0.1)[26] and tidyr (v 0.8.3)[27] in scripts as linked throughout the methods. Statistical analyses were carried out in R, as detailed in the results section. The sign test was carried out using the R package BSDA (v 1.2.0) [28].

**Figure 3:**
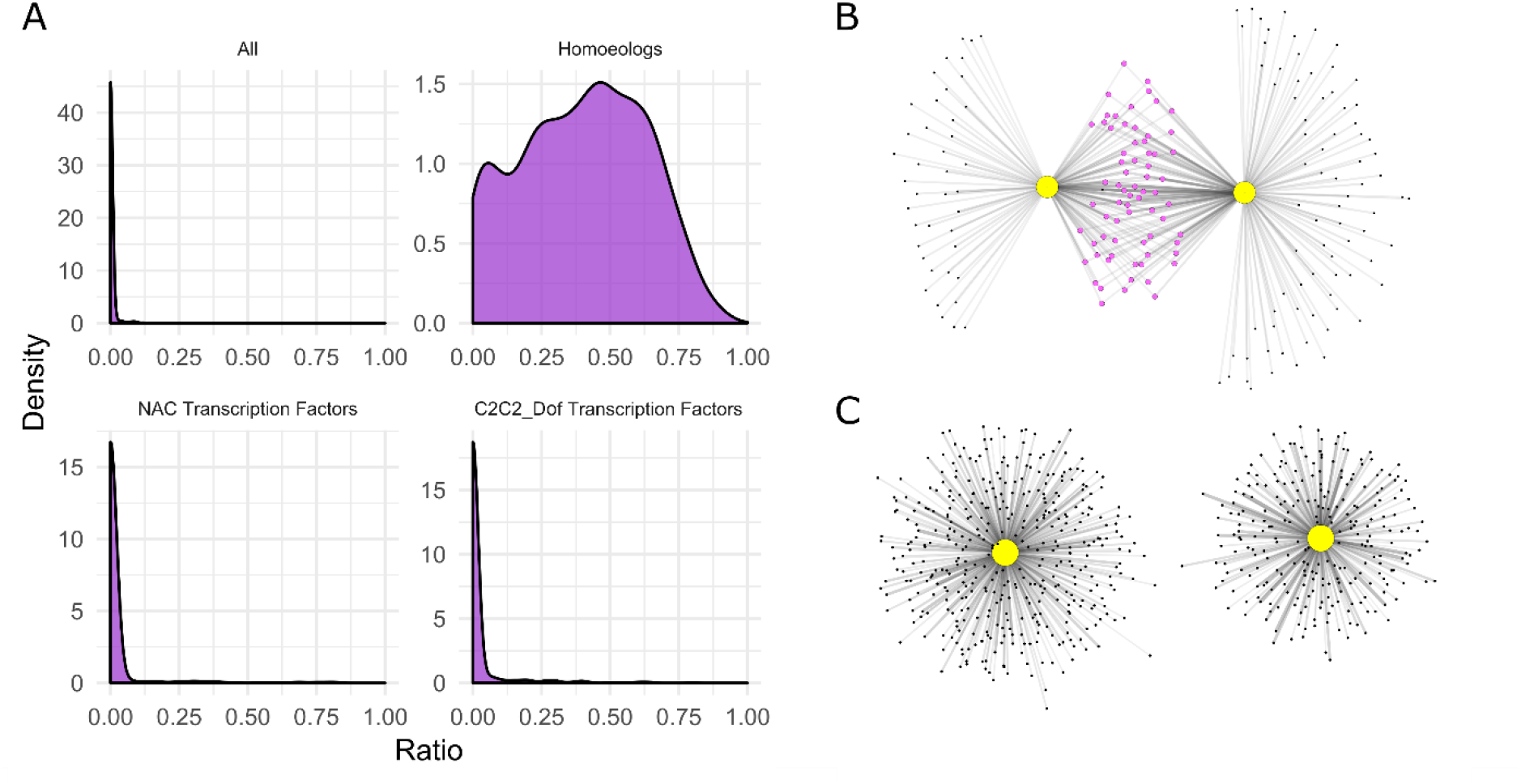
Non-homoeologous transcription factors share few targets in the GENIE3 network. (**A**) Comparison of the shared targets of 1000 random transcription factor pairs found that the majority of transcription factors share few to no targets in common. This was also found to be the case within the majority of transcription factor families, showing here NAC and C2C2_Dof transcription factors. However, pairs of homoeologous transcription factors shared many more targets in common, with a mean shared ratio of 39.7%. (**B**) An example of shared overlap between two homoeologous transcription factors, the NAC transcription factors *NAM-A1* (TraesCS6A01G108300) and *NAM-D1* (TraesCS6D01G096300) is shown, with the two homoeologs sharing 62% of possible targets. (**C**) The two randomly chosen transcription factors, in this case TraesCS1D01G436500 (Sigma 70-like family) and TraesCS4B01G383400 (HSF family), share no targets.

## Results

### RNA-Seq analysis

In 2014, Pearce *et al.* analysed the differences in gene expression between wild type (WT) Kronos, a tetraploid wheat cultivar, and a *NAM-A1* loss-of-function mutant (*nam-a1* or *gpc-a1*) which contained a premature stop codon (W114*) [14]. Here, we reanalysed the RNA-Seq datasets for the wild-type and *nam-a1* single mutant lines at 12 and 22 DAA using the most recent wheat genome annotation [1]. Reads were pseudoaligned to the A and B genomes of the RefSeqv1.1 transcriptome using kallisto [17], a software which has been shown to differentiate well between homoeologs during alignment and is thus appropriate for use with polyploid wheat [2, 8]. Each sample contained on average 35 million reads, with the exception of one sample with 85 million reads, of which on average 78% were aligned to the transcriptome (Supplementary Table 1).

To identify genes differentially expressed between the WT and *nam-a1* mutant at the two developmental timepoints (12 and 22 DAA) we used sleuth, a program designed for analysis of RNA-Seq experiments for which transcript abundances have been quantified with kallisto [18]. Using a relatively relaxed cut-off of *q* < 0.05, we identified 866 differentially expressed genes (DEGs) between WT and the *nam-a1* mutant at 12 DAA and 130 DEG at 22 DAA (Supplementary Table 2). This set of DEGs will be referred to as the *de novo* DEGs throughout the paper. We carried out gene ontology (GO) term enrichment analysis on the two sets of DEGs and found that DEGs at both timepoints are highly enriched for terms related to metal ion transport, including zinc, manganese, and copper (p < 0.001, adjusted for false discovery rate; Supplementary File 2). This correlates closely with the findings from the original analysis, which found that GO terms related to transporter function were highly regulated by *NAM-A1* [14]. This also supports previous physiological studies of the *NAM* genes which found them to be important in nutrient remobilisation and transport [15, 29].

We used BLASTn to compare the differentially expressed sequences from the Pearce dataset to the RefSeqv1.1 transcriptome annotation which was used for the *de novo* analysis [1]. We successfully assigned 453 out of 517 DEG identified by Pearce *et al.* to RefSeqv1.1 gene models (442/504 DEG at 12 DAA and 11/13 DEG 22 DAA). Based on this common nomenclature, we then directly compared the *de novo* DEGs identified by sleuth with the DEGs identified originally [14]. At 12 DAA, 177 of the 442 DEGs (40%) were present in the *de novo* differential expression set, whereas at 22 DAA, 7 of the 11 DEGs (64%) were found in the *de novo* set.

At both developmental time points, our reanalysis identified more transcripts as significantly differential expressed compared to the original Pearce *et al.* study. This may be due to the more liberal significance cut off used (*q* < 0.05) in the current analysis and/or the fact that a combination of four statistical tests were used to reduce false positive discoveries in the original study [14]. To determine the impact of the cut-off value on the calling of DEGs, we ranked the *de novo* DEGs by *q*-value and recorded the position of the 177 shared DEGs at 12 DAA. We found that the majority of shared DEGs (53%) ranked in the top quarter of the list of *de novo* DEGs (Supplementary Figure 1). However, 9% of the common DEGs were found in the bottom quarter of the *de novo* DEGs. This suggests that the cut-off value of *q* < 0.05 was an appropriate level to maximise the identification of relevant DEGs.

### The GENIE3 network predictions overlap with known DEGs

To investigate whether the GENIE3 network provides biologically relevant information, we compared the GENIE3 predicted targets for *NAM-A1* against the list of differentially expressed genes identified between the wild-type and *nam-a1* mutants from the *de novo* RNA-Seq analysis. As the RNA-Seq experiment was carried out in tetraploid wheat, we only considered target genes on the A or B genome. We focussed on the 12 DAA and 22 DAA timepoints, which captured the onset and intermediate stages of senescence, respectively. At 12 DAA, we found that of the 79 genes predicted to be targets of *NAM-A1* in the GENIE3 network, 12 were shared with the set of *de novo* DEGs (15.2%; Figure 1A). However, at 22 DAA only 4 of the 79 GENIE3 predicted targets were shared with the DEGs (5.1%; Figure 1A). The decrease in overlap between 12 and 22 DAA is consistent with *NAM-A1* primarily acting early in senescence [14, 15].

We then compared the lists of DEGs at 12 and 22 DAA against the targets of all 3,384 transcription factors included in the GENIE3 dataset (Figure 1B). The median number of shared targets between the DEGs and predicted targets of a given transcription factor was 0, with a maximum of 33.3%. Comparing the overlap between random transcription factors and the RNA-Seq dataset, we found a significantly higher level of overlap between the GENIE3-predicted targets of *NAM-A1* and genes differentially expressed in the *nam-a1* mutant at both timepoints (p < 2.2e-16, Sign Test; Figure 1B). This result suggests that the GENIE3 network has value in directing focus towards targets with independent experimental support.

### Identification of senescence associated transcription factors

We hypothesised that transcription factors which also share predicted targets with the *de novo* DEGs may have roles in senescence. We therefore identified transcription factors that had a higher percentage of shared targets with the *de novo* DEGs than *NAM-A1* itself (Figure 2C, Supplementary Table 3). In total, we identified 20 such transcription factors, 0.6% of all transcription factors in the network. Five transcription factors were identified through comparison to the 12 DAA DEGs, 11 with the 22 DAA DEGs, and a further four which had a higher shared ratio at both time points. We obtained expression data for all 20 of the transcription factors across a developmental time-course and a senescence-specific time-course [2, 3]. A diverse range of transcription factor families were represented, including four NACs, from the same family as *NAM-A1* (Supplementary Table 3). Only one pair of homoeologs was identified, from the C2C2-CO-like family. Using the developmental time-course, we calculated the fold-change in gene expression between non-senescing tissues and senescing leaf tissues as previously described [3]. We found that 14 of the 20 genes showed an increase in expression in the senescing tissues, four of which were enriched more than two-fold (Figure 2A). Based on a published analysis of the senescence-specific time-course, six of the 20 transcription factors were identified as differentially regulated during flag leaf senescence in wheat [3]. Analysis of the GENIE3 predicted targets for these 20 genes identified nine transcription factors which shared at least one target gene with *NAM-A1* (Figure 2B).

To investigate the potential functions of these 20 transcription factors further, we identified GO terms which were enriched in the GENIE3-predicted targets of these transcription factors (Supplementary Table 3; Supplementary File 2). At 12 DAA, the targets of three of the nine transcription factors were enriched for transporter-related GO terms, while others were enriched for senescence-related GO terms such as catabolism, phosphatase activity, and chlorophyll biosynthesis. Of the transcription factors identified at 22 DAA, GO terms related to transporters, senescence, circadian rhythms, and stress responses were enriched in the GENIE3-predicted targets. By integrating the information from the GENIE3 network with independent senescence-related expression data, we were able to identify a robust set of candidate senescence-associated transcription factors to prioritise for functional characterisation.

### Non-homoeologous transcription factors share few targets in the GENIE3 network

After establishing that the GENIE3 network can provide biologically-relevant predictions, we then turned to using the network to interrogate genome-wide patterns in transcription factor targets. We first investigated the extent to which transcription factors share the same targets. To do this, we carried out pairwise comparisons between randomly selected transcription factors and calculated the overlap between their targets. Following 1,000 iterations, we found that any two random transcription factors typically have no targets in common, with a median and 3^rd^ quartile shared ratio of 0%. The distribution was highly positively skewed, as the vast majority of comparisons shared 0 targets (Figure 3A, B; Table 1). However, there was a long tail to the right of the graph, highlighting that in some cases, certain transcription factors do share a substantial portion of targets.

**Table 1:**
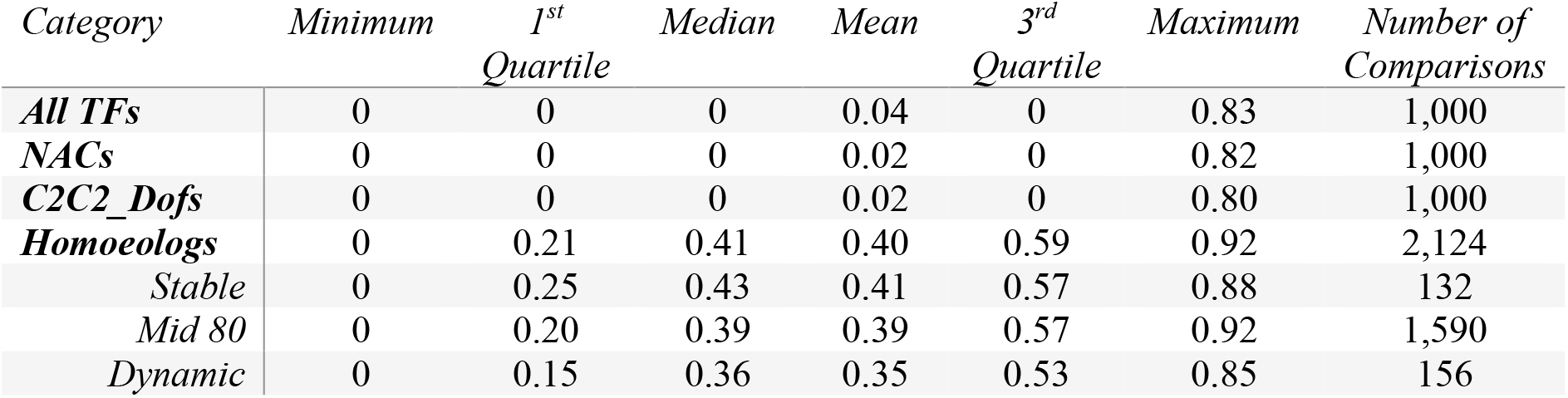
Summary statistics for the shared ratio distributions.

The set of transcription factors in the GENIE3 network was then split into separate transcription factor super-families, as previously annotated [1]. The same analysis was performed within each transcription factor family, carrying out pairwise comparisons between transcription factor targets. We found that for the majority of transcription factor families, the distributions of shared targets were very similar to that found for the full set, as illustrated by the representative NAC and C2C2-Dof families (Figure 3A, Table 1; Supplementary Figure 2). Not all transcription factor families were large enough to support 1,000 unique pairwise comparisons, and in these cases the distribution clearly deviates from the whole (e.g. in CCAAT_HAP3, N = 3).

Roughly 70% of wheat homoeolog triads (composed of A, B, and D genome copies) show balanced expression patterns, that is, similar relative abundance of transcripts from the three homoeologs across tissues [2]. We therefore hypothesized that homoeologs would be more likely to share predicted targets than randomly selected transcription factors, even within the same family. To test this, we randomly selected syntenic triads from the GENIE3 dataset, and calculated the percentage of shared genes for each of the three pairwise comparisons (A-B, A-D, B-D). This was carried out for the 708 syntenic triads included in the network, a total of 2,124 pairwise comparisons, and showed that homoeologs are indeed more likely to share targets with each other than randomly selected triads are (Figure 3A, C, Table 1).

### Dynamic triads share fewer targets than stable triads

Wheat genome triads can be classified into different categories based on how the expression levels of the homoeologs varies across tissues [2]. So-called ‘dynamic’ triads represent the 10% of triads which show the highest variation in relative expression levels of the different homoeologs across tissues, while ‘stable’ triads represent the 10% of triads with the lowest variation. Dynamic triads are under more relaxed selection pressure and we hypothesised that they may represent the initial steps toward neo or sub-functionalization of wheat homoeologs [2]. This hypothesis would suggest that homoeologs in dynamic triads are more likely to have distinct functions, and thus may be less likely to share predicted targets. To test this, we compared the level of overlap between targets of homoeologous transcription factors in dynamic and stable triads, as well as the ‘Mid 80’ intermediate set. We found that the dynamic triads do indeed have significantly less overlap in targets than the stable triads (p < 0.05, Wilcox test; Figure 4A; Table 1), supporting the neo/sub-functionalization hypothesis.

**Figure 4:**
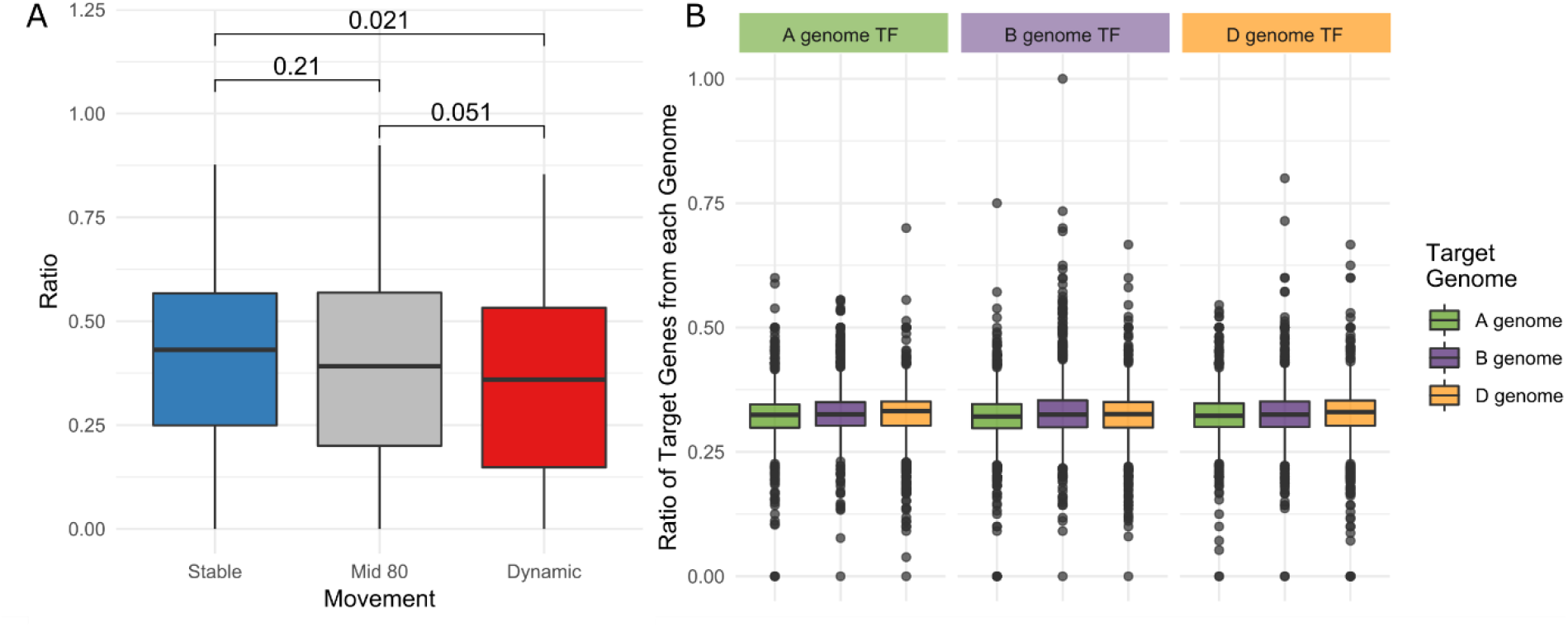
The effect of relative homoeolog expression variation on shared ratios. (**A**) There is a significant reduction in shared targets between homoeologs in dynamic triads compared to stable triads (Wilcox test, p = 0.021). There is a near-significant decrease between the Mid 80 subset and the Dynamic triads (p = 0.051), and a slight non-significant decrease from Stable to Mid 80 triads (p = 0.21). (**B**) We found no evidence that transcription factors from a specific genome were more or less likely to have targets from the same genome (Two-sample Kolmogorov Smirnov test, FDR adjusted).

We next examined whether the targets of a transcription factor may hold signatures of the evolutionary origin of that transcription factor. We hypothesized that a transcription factor is more likely to have targets that reside on the same genome as the transcription factor itself; *e.g.* GENIE3-predicted targets for an A genome transcription factor will be enriched for A genome targets compared to B and D genome targets. We found no significant association between the genome of origin of the transcription factor and the genomes of its targets (Figure 4B). Likewise, we found no significant association between transcription factor genome and target genome when triads were assigned into their respective movement categories. For example, an A genome transcription factor from a Dynamic triad had similar numbers of GENIE3-predicted targets from each of the three genomes. (Supplementary Figure 3).

### GO-term enrichment of predicted targets suggests novel biological functions

We next examined if we could use the GENIE3-predicted targets to gain a more general overview of functional trends within transcription factor families. Using the GO terms described in [2], we carried out a GO-term enrichment on the targets of individual transcription factor families (Supplementary File 2). To test the validity of this method, we first checked the predicted GO terms of the Heat Stress Factor (HSF) transcription factor family. Based on their known role in responding to heat and light stress [30], we expected to see GO terms highly enriched for heat and light stress responses. This was the case, suggesting that this approach can provide accurate insights into the large-scale functions of transcription factor families.

We then identified enriched GO terms for the remaining 56 transcription factor families. Some families were enriched for few or no GO terms, and we found that this was typically the case for families with few members. We restricted our analysis, therefore, to the 34 transcription factor families with at least 30 members. In many cases, enriched GO terms supported known functions of transcription factor families. For example, the MADS_II family was enriched for, amongst other terms, floral organ identity (GO:0010093) which corresponds to their known role in flower patterning in wheat and other species [31]. The mTERF family was strongly enriched for chloroplast-related terms, corroborating their known role in organelle function [32, 33] while the WRKY family was highly enriched for stress-response GO terms [34-36].

Beyond the expected enriched GO terms, we also identified some cases where highly significantly enriched GO terms may point towards a previously unknown function within that transcription factor family. For example, one of the more highly enriched GO terms for the C2C2-Dof family is positive gravitropism (GO:0009958). This, combined with other GO terms related to amyloplasts and auxin responses, suggests that members of the C2C2-Dof family may play a role in regulating the gravitropic growth of roots [37]. We also see that the TUB family, of Tubby-like transcription factors, is highly enriched in cell-cycle related GO terms. This includes specific terms involved in microtubule movement and spindle formation (GO:0007018 and GO:0051225) as well as in regulation of cell cycle progression and transition (GO:0010389, GO:0051726 and GO:0000911). Plant TUB proteins contain an F-box domain, suggesting they may function in tandem with other F-box-containing proteins, such as SCF E3 ubiquitin ligases, which also regulate the cell cycle in plants [38].

### The GENIE3 network is accessible through KnetMiner

We have made the GENIE3 network available in the KnetMiner discovery platform for wheat (https://knetminer.rothamsted.ac.uk/) [39]. The top one million TF-target predictions were integrated onto the wheat genome-scale knowledge graph [16] as directed relations between genes (A regulates B). The interaction weight for each predicted relationship was also included in the network, where larger weights correspond to more strongly supported relationships. The data is accessible in conjunction with other information types (e.g. protein-protein interactions, literature co-occurrence, ontologies, homology, etc.) and can be searched through the KnetMiner web-app or web-services.

KnetMiner can be searched using keywords, genes or genomic regions to identify connections between genes and hidden links to complex traits. Searching for “NAM-A1” returns two hits, TRAESCS6A02G108300 and TRAESCS6D02G096300, which correspond to *NAM-A1* and its D-genome homoeolog *NAM-D1*, respectively. Using the KnetMiner network, we can visualise the relationships between *NAM-A1* and *NAM-D1*, seeing, for example, that they target each other in the GENIE3 network (Figure 5). Associated traits are also included in the network, linking *NAM-A1* and *NAM-D1* to “Grain Protein Content” (Figure 5). Links to orthologous genes in other species are also included in the network, such as the *Arabidopsis thaliana* orthologue *ANAC018* (Figure 5).

**Figure 5:**
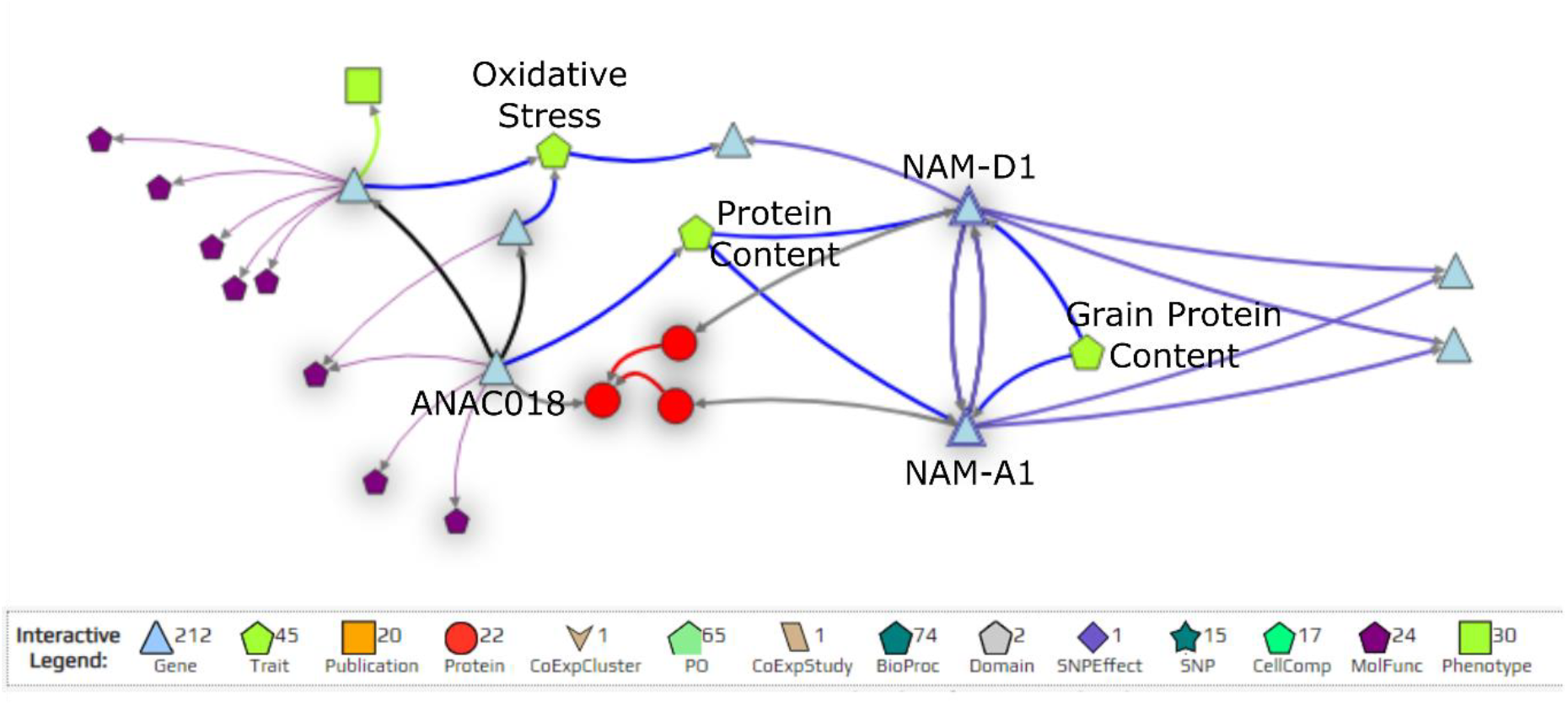
The KnetMiner network depicts connections with *NAM-A1*. The wheat transcription factors *NAM-A1* and its homoeolog *NAM-D1* target each other in the GENIE3 network (purple arrows) and share some target genes (blue triangles) in common. The network also identifies traits associated to genes (green pentagons), such as “Protein Content” for *NAM-A1, NAM-D1*, and the *A. thaliana* orthologue *ANAC018.* The legend below the network describes the meaning of each shape in the network. Not all connections present in the KnetMiner network are depicted in the figure; only a subset are shown for clarity.

## Discussion

Here we have shown that a GENIE3 network provides biologically-informative predictions of targets for transcription factors in polyploid wheat. We have used the network in conjunction with independent RNA-Seq datasets to identify a set of candidate senescence regulators. We have also shown the value of the network to analyse genome-wide patterns of homoeologous transcription factors and transcription factor families.

### Re-analysis of an RNA-Seq dataset identifies a high-quality set of differentially expressed genes

In our analysis of the GENIE3 network, we used a previously published RNA-Seq dataset to validate its ability to predict the targets of a well characterised transcription factor, *NAM-A1* [15]. In doing so, we needed to re-align the raw RNA-Seq reads to the most current transcriptome, RefSeqv1.1 [1]. After carrying out *de novo* pseudoalignment and differential expression analysis, we obtained a larger set of differentially expressed genes between the wild-type and mutant *nam-a1* lines than the original study [14]. In part, this is likely due to the less strict cut-off used in our study (q < 0.05). In the original study [14], a combination of four statistical tests were used to reduce the number false positive discoveries. However, as our intention was not to reduce the rate of false positives, but rather that of false-negatives or incorrectly removed genes, we used a less stringent cut-off, based solely on adjusting the original *q*-values for false discovery rate. We found that the DEGs identified in Pearce *et al.* [14] were found throughout the *de novo* list of DEGs, suggesting that the cut off chosen was not overly generous (Supplementary Figure 1). We also closely recapitulated their findings in that our *de novo* DEGs were highly enriched for metal ion transport-related GO terms, indicating that the *de novo* DEGs are consistent with *NAM-A1* function. These enrichment results also corroborate our understanding of *NAM-A1* as a critical regulator of nutrient remobilisation during senescence.

### The GENIE3 network provides biologically relevant transcription factor - target relationships

Gene networks are increasingly used in plant genetics research as a way to establish relationships across large gene sets and for hypothesis-generation. Initial assessment of the biological relevance of these gene networks often rely on enrichment analyses using GO terms. These methods are useful in identifying trends within large gene sets, as we found when carrying out GO term enrichment of transcription factor families. The targets of families with known functions, such as the Heat Stress Factors, were enriched for the expected GO terms [30]. However, these enrichment analyses are limited by the information that is used to develop the GO terms themselves. Very few GO term annotations are supported by experimental evidence even in model species such as *Arabidopsis thaliana* [40]. GO terms associated to the RefSeqv1.0 transcriptome were based primarily on automated annotation and orthologues in other plant species, with over 96% of the GO terms assigned to genes based on inference from sequence orthology (ISO) [1]. As a result, while enrichment analyses with GO terms can provide useful information particularly with large-scale datasets, they must be combined with external data to validate their predictions.

As a result, while the GO term enrichment analyses suggested that the network produced biologically relevant results, validation of the network required the use of experimentally-derived data. By using independent RNA-Seq datasets, which were not used in the creation of the GENIE3 network, we were able to show that the predictions made by the *in silico* network hold up under comparison to *in vivo* datasets. It is important to note, however, that the overlap between the predicted targets of *NAM-A1* and the differentially expressed genes from the RNA-Seq data is far from complete. At 12 DAA, the timepoint where *NAM-A1* is thought to first start exerting its effect, only 12 genes are shared out of 79 predicted genes and 866 differentially expressed genes. While this gives a shared ratio of approximately 15%, significantly higher than that expected by chance, the GENIE3 network and the differentially expressed genes do not contain identical targets. However, based on the differences in the datasets used, it is likely that a large portion of this discrepancy is due to the fact that the GENIE3 network was derived from 850 distinct RNA-Seq samples, spanning different tissues, ages, stresses, and varieties, while the RNA-Seq dataset came from single timepoints taken from flag leaf tissue [1, 14]. While *NAM-A1* is expressed in the flag leaf during senescence, it is also expressed in the peduncle during senescence [2, 8], and at lower levels in various leaf, stem, and even spike tissues during development (Supplementary Figure 4). It’s possible that many, if not most, of the predicted targets from the GENIE3 network may be regulated or influenced in some way by *NAM-A1*, but not in the flag leaf and at the precise 12 DAA and 22 DAA timepoints captured in the independent RNA-Seq data.

### The GENIE3 network can be used for hypothesis generation with large gene sets

The fact that any two random transcription factors share, on average, zero targets in common in the GENIE3 network highlights that the network is not overwhelmed by spurious connections between transcription factors and biologically irrelevant targets. The network is also not so conservative that transcription factors with similar functions share no targets in common, as is made clear by the more normally-distributed homoeologs (Figure 3A). Nevertheless, as most transcription factors do not share targets, this suggests that the original cut-off chosen for the network (1,000,000 connections) was appropriately stringent to avoid noisy, low-confidence connections.

The presence of overlapping targets between transcription factors suggests that the transcription factors may be acting in similar pathways. However, it is important to recognise the limitations of the network, in particular that predicted targets of a transcription factor are not necessarily true targets. Therefore, integrating the GENIE3 network with other sources of regulatory information, such as RNA-Seq datasets as demonstrated here, can provide cumulative evidence towards new hypotheses and candidate genes. By combining the DEGs obtained from a *nam-a1* mutant line and the GENIE3 network, we have produced a set of candidate transcription factors which may act in the same or similar pathways as *NAM-A1*. We have then compared these candidates with an independent senescence time-course [3], highlighting six candidate genes which were identified through both methods and are good candidates for further exploration. Moving forwards, information generated from networks such as the GENIE3 will need to be validated functionally and *in planta*. Recently, *NAM-A2*, a previously uncharacterised member of the *NAM* family of transcription factors, was predicted to be involved in senescence based on expression and network data [3]. The function of this transcription factor was then validated using independent mutant TILLING lines [41] demonstrating the ability of the networks to predict biologically-relevant candidates.

### Homoeolog expression variation in dynamic triads may be indicative of functional divergence

Previous work showed that triads of homoeologs can display variable patterns of genome dominance across different tissues [2]. Triads with the most variable relative homoeolog expression patterns, ‘Dynamic’ triads, were also found to have higher Ka/Ks values, suggesting they were under reduced selection pressure. It was proposed that the variation in relative expression patterns across tissues arises as a result of this relaxed selection pressure, facilitating both neo- and sub-functionalisation following polyploidy. We found that dynamic triads were less likely to share GENIE3 targets in common than stable triads, supporting the hypothesis that dynamic triads are in the process of diverging functionally (Figure 4B). However, we found no correlation between the genome of origin of the transcription factor and the target genomes in dynamic triads (Supplementary Figure 3).

At what stage, then, did the targets of homoeologs in dynamic triads begin to diverge? These results may suggest that the variation in expression seen between the homoeologs arose following polyploidisation, as there is no bias towards the genome of origin. However, we do not know enough about the behaviour of transcription factors following polyploidisation to draw clear conclusions. For example, we do not know to what extent transcription factors gain the ability to regulate homoeologous genes on other genomes after hybridisation. The application of methods such as ChIP-Seq [42], DAP-Seq [43], and large-scale yeast two-hybrid interaction screens [44] to transcription factors in diploid and polyploid wheat will provide experimental data on homoeologous transcription factor interactions and binding preferences. Until these datasets become available it is premature to draw conclusions on the evolutionary origins of transcription factor homoeolog functional divergence. Nevertheless, genome-wide analyses of datasets such as the GENIE3 network and the expression datasets on expVIP have pointed to the dynamic triads as a source of genetic functional variation [2, 8].

## Conclusions

Using publicly available datasets, we have shown that the wheat GENIE3 network provides biologically-relevant information that can be used to guide hypothesis generation in wheat. The utility of the network lies particularly in enrichment analyses of larger gene sets and in integration with other datasets, such as independent RNA-Seq experiments, for candidate gene selection. New germplasm resources in wheat, such as the *in silico* TILLING resource [41], can be rapidly leveraged for functional characterisation of candidate genes *in planta*. Transgenic approaches such as CRISPR [45] and virus-induced gene silencing (VIGS) [46] can now be used in wheat to validate gene function. The GENIE3 network can be accessed through the KnetMiner web application and using R scripts available from https://github.com/Uauy-Lab/GENIE3_scripts. We predict that gene regulatory networks such as GENIE3 will play an increasingly important role in wheat genetics as more transcriptomic datasets become publicly available.

## Acknowledgements

We would like to acknowledge A. Braeutigam (Bielefeld University) for her useful comments during the development of the manuscript, and for her work developing the GENIE3 network.

## Author Contributions

SAH and CU conceived the study; SAH and AB carried out the RNA-Seq analysis and validation; SAH carried out the genome-wide analyses of the network; AS and KH-P integrated the GENIE3 dataset into KnetMiner; SAH wrote the manuscript, with contributions from CU, AB and KH-P. All authors have read and approved the final manuscript.

## Data Availability

All scripts used for analysis in this paper are available on Github at https://github.com/Uauy-Lab/GENIE3_scripts. Links to the public datasets used in the analysis are included within the Materials and Methods, where appropriate, or are linked in the scripts on Github. In general, datasets from the Ramírez-González *et al.* 2018 paper are available at https://opendata.earlham.ac.uk/wheat/under_license/toronto/Ramirez-Gonzalez_etal_2018-06025-Transcriptome-Landscape/. The original GENIE3 network is deposited at https://doi.org/10.5447/ipk/2018/7.

## Supplementary Files

Supplementary Table 1: Kallisto mapping statistics.

Supplementary Table 2: List of *de novo* differentially-expressed genes.

Supplementary Table 3: List of candidate senescence regulators, as shown in Figure 2.

Supplementary File 1: Contains Supplementary Figures 1-4.

Supplementary File 2: Contains the enriched GO terms for the GENIE3 targets of all transcription factor families and of the candidate senescence regulators (Supplementary Table 3). See README file in folder for further details.

