## Supplementary File 1 for "Validation and characterisation of a wheat GENIE3 network using an independent RNA-Seq dataset"

Supplementary Information.

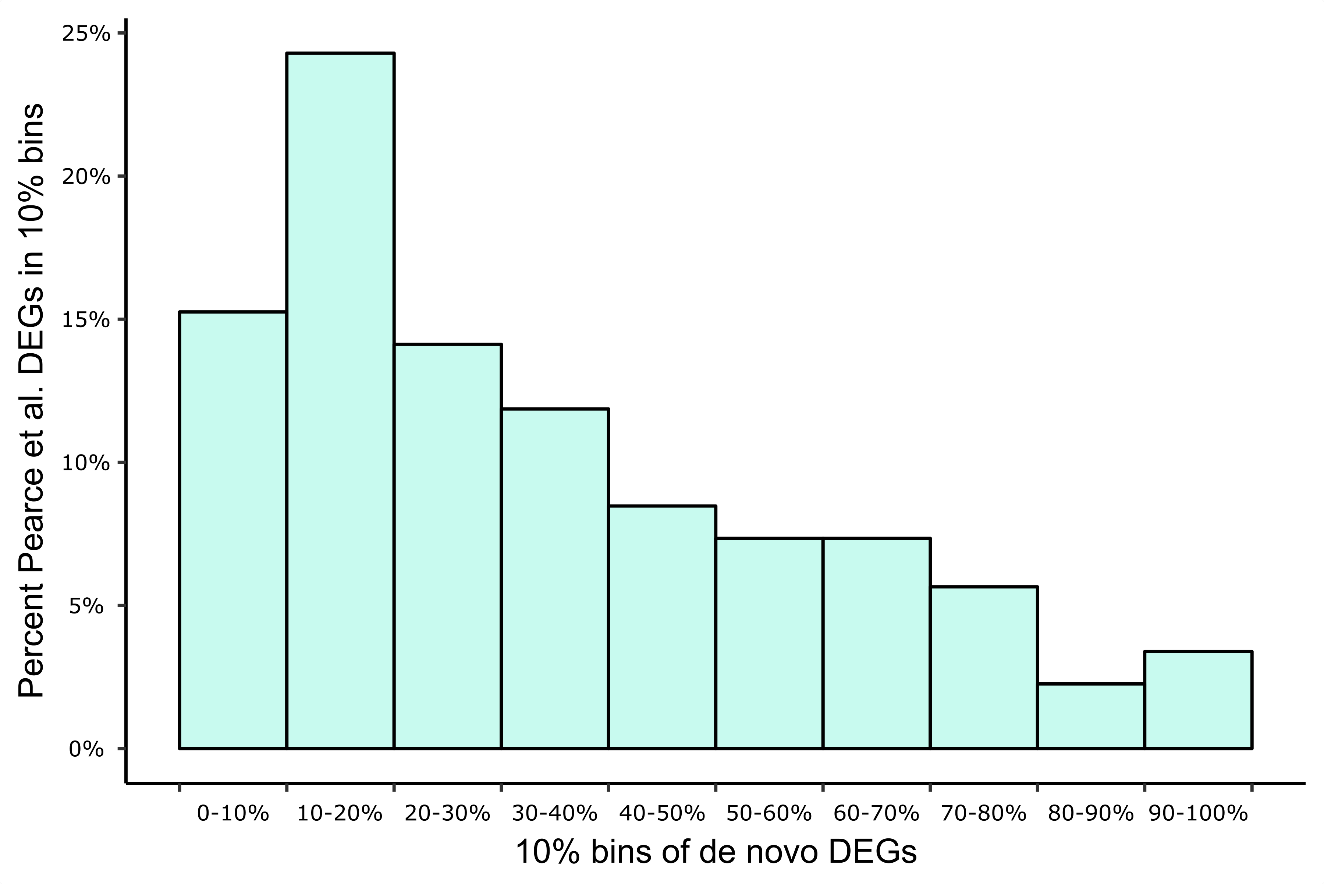

**Supplementary Figure 1: Differentially expressed genes from Pearce *et al.* 2014 are present in the *de novo* gene set.** The majority (53%) of differentially expressed genes (DEG) from Pearce et al. fall into the top 25% of the *de novo* DEG set [14]. 9% of the Pearce *et al.* DEGs fall into the bottom 25% of the *de novo* DEGs. DEGs are ranked based on q-value, where the lowest (most significant) q-value is ranked at the top.

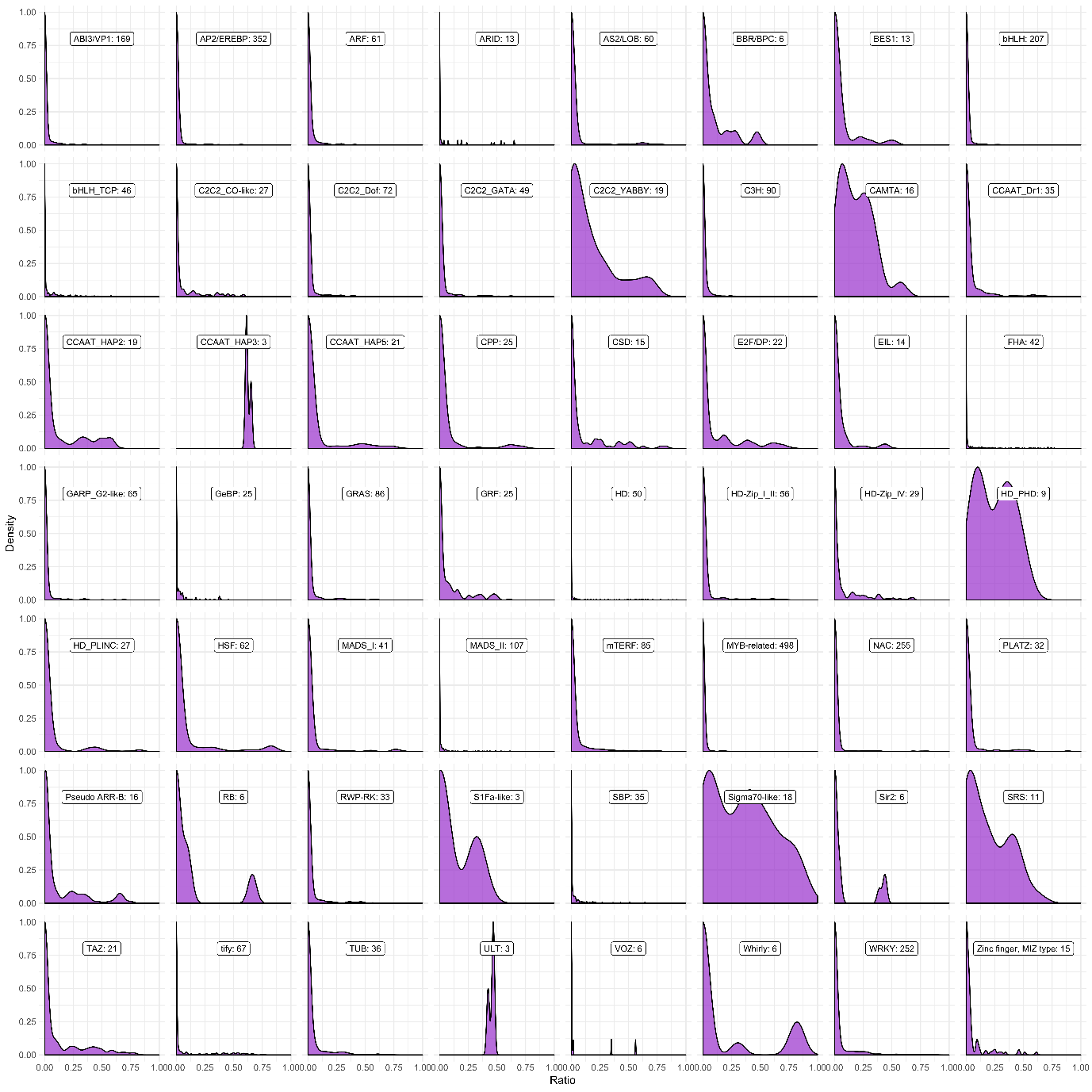

**Supplementary Figure 2: Shared downstream targets between transcription factors in the same superfamily.** Density distributions for the shared ratios of unique transcription factor pairs sampled from each of the 56 superfamilies tested closely recapitulate the density distribution seen for all random transcription factors (Figure 3A). Superfamilies with few members are less skewed right, due to the increased proportion of intra-triad comparisons included in the distribution.

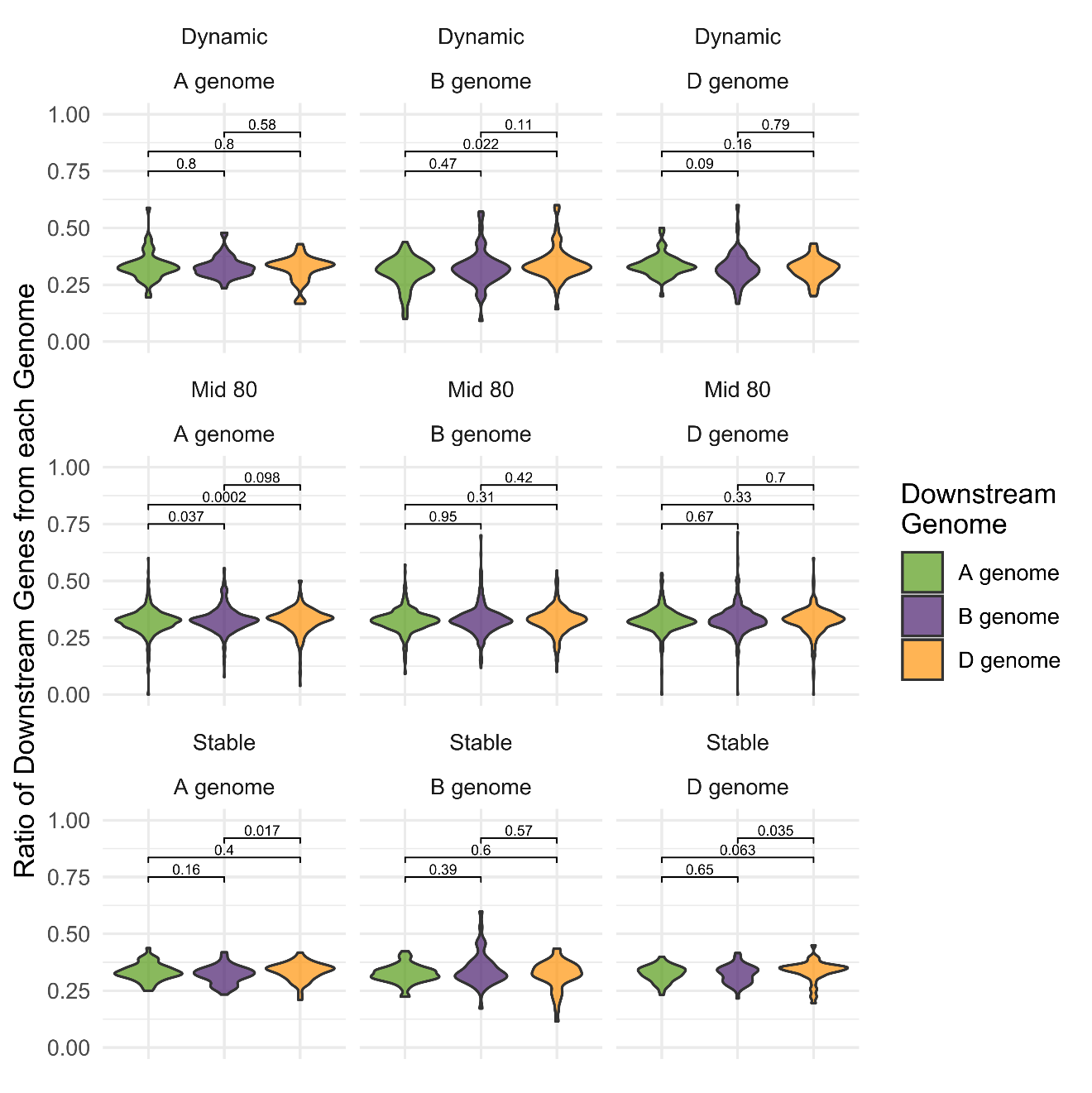

**Supplementary Figure 3: No clear pattern is seen between transcription factor movement category and genome bias in downstream genes.** Percentage of downstream genes from each genome is shown for all transcription factors, split by transcription factor genome and movement category. The *p*-value for each pairwise test between distributions is shown in each panel (Wilcox test).

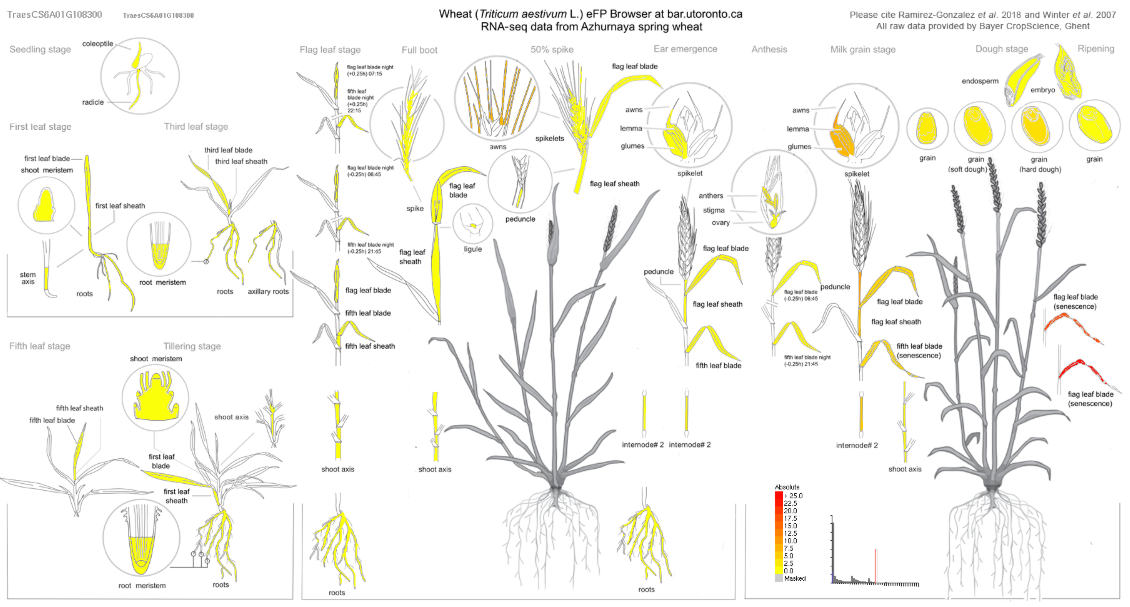

**Supplementary Figure 4: *NAM-A1* is expressed predominantly in the senescing flag leaf.** Expression data from the developmental time course of the spring wheat variety Azhurnaya is displayed using the Wheat eFP browser (<http://bar.utoronto.ca/efp_wheat/cgi-bin/efpWeb.cgi>) [2]. *NAM-A1* is predominantly expressed in the senescing flag leaf (up to approximately 50 TPM), but also in the peduncle (9.8 TPM) and the glumes (7.8 TPM) among other tissues.
