## Supplementary File 2 for "Validation and characterisation of a wheat GENIE3 network using an independent RNA-Seq dataset": README.txt

README for Supplementary File 2

This file contains the Gene Ontology (GO) enrichment for the GENIE3 targets of different genes or sets of genes.
GO terms were filtered by over-represented p-value (adjusted for false discovery rate), only retaining those with p < 0.05
"numDEinCat" gives the number of genes in the set of targets that are associated with this GO term, "numinCat" is the number of genes in the universe that are associated with this GO term.

Subfolders:

-Higher_Ratio_Genes
This folder contains the GO term enrichment for each of the 20 candidate senescence regulators listed in Supplementary Table 3 and Figure 2. 
GO term enrichment was carried out on the individual targets of each transcription factor (TF); note that some TFs have few targets; in these cases the GO term enrichment is unlikely to be informative.

-TF-Families
This folder contains the GO term enrichment for each of the transcription factor families, 57 in total. 
GO term enrichment was carried out on the individual targets of each transcription factor that is part of that family. 
I.e. enrichment was carried out on the targets of all NAC transcription factors.
